# Electrophysiology prediction of single neurons based on their morphology

**DOI:** 10.1101/2020.02.04.933697

**Authors:** Mario Michiels

## Abstract

Electrophysiology data acquisition of single neurons represents a key factor for the understanding of neuronal dynamics. However, the traditional method to acquire this data is through patch-clamp technology, which presents serious scalability flaws due to its slowness and complexity to record at fine-grained spatial precision (dendrites and axon).

In silico biophysical models are therefore created for simulating hundreds of experiments that would be impractical to recreate in vitro. The optimal way to create these models is based on the knowledge of the morphological and electrical features for each neuron. Since large-scale data acquisition is often unfeasible for electrical data, previous expert knowledge can be used but it must have an acceptable degree of similarity with the type of neurons that we are trying to model.

Here, we present a data-driven machine learning approach to predict the electrophysiological features of single neurons in case of only having their morphology available. To solve this multi-output regression problem, we use an artificial neural network that has the particularity of providing a probability distribution for every output feature, to incorporate uncertainty. Input data to train the model is obtained from from the Allen Cell Types database. The electrical properties can depend on the morphology, whose acquisition technology is highly automated and scalable so there exist large data sets of them. We also provide integrations with the BluePyOpt library to create a biophysical model using the original morphology and the predicted electrical features. Finally, we connect the resulting biophysical model with the Geppetto UI software to run all the simulations in a sophisticated user interface.

## 1 Introduction

The brain physiology consists of many interconnected components that need to be studied at multiple scales (Sejnowski et al., 2014). We can order these scales based on their physical size. At the microscopic level, we have the genomics scale that takes place in the cell core. The next level is the cellular one, which involves the study of morphology and electrophysiology of single cells. At this level, the common acquisition technologies comprehend multiple kinds of microscopy, single cells recording via patch-clamp, etc. Finally, at the upper level we study groups of cells categorizing the brain in multiple lobes/-parts. Here we can use non-invasive recording techniques like EEG, fMRI, etc. This paper focuses on the study of single cells, thus it only takes place at the cellular level.

The electrophysiology of single neurons has been widely studied since decades to understand how the brain works from the cellular perspective. One specific purpose of electrophysiological measurements is the creation of biophysical detailed models to simulate experiments in silico (Markram, 2006). These models are particularly useful due to the difficulty and cost to obtain in vitro measurements. Patch-clamp is the most common method to record electrical signals from single neurons, which requires to inject a small current into the soma in order to stimulate it and record the resulting response.

Patch-clamp recording, while precise, has two main drawbacks: slowness and complexity to record at detailed spatial precision (dendrites and axon). The first is because it is inherently designed to inject and record from single neurons. However, its speed has been nowadays increased by robotic technology to manage multiple pipettes at once (Annecchino and Schultz, 2018), although it is still not enough to record hundreds of neurons of the same tissue at once. The spatial precision otherwise is due to the difficulty to inject and record from the dendrites or axons since they are very thin and the pipette can not properly penetrate without damaging the cell. It is still a feasible technique and some laboratories were able to do patch-clamp in the dendrites (Larkum et al., 2001) but it is so complex that is not a scalable technique to do it in many cells.

It is well known that the electrophysiology depends on morphological and molecular features. Morphologies have been studied since the Ramón y Cajal era (Sotelo, 2003) and they have been categorized in multiple classes that allow us to define a specific behavior for each one. Lastly, the molecular profile regulates the ion channels kinetics. Therefore, in theory, the electrical properties can be defined by the location of the ion channels in their morphology and their kinetics regulated by the molecular profile (Toledo-Rodriguez et al., 2004). However, in practice all the relationships between the morphology, molecular and electrophysiological features are not known yet.

Markram et al. (2015) proposed to classify the neurons as a morphological type and a electrical type, thus creating a morpho-electrical pair that defines the association. This is a general high level approach that can not distinguish between different neurons in the same class and can not fully comprehend neurons that would not fit perfectly in a specifically defined class.

Currently, morphology data acquisition can be fast and automatized via image detection from microscopy data. There also exist large databases like NeuroMorpho.org (Ascoli et al., 2007) to store them and tools like NeuroSuites (Michiels, 2019) to easily analyze them. Molecular data can also be collected in a fast way with technologies like microarrays but the analysis becomes more complex as there can be many factors that affect the dynamics of the molecular behavior. On the other hand, electrical patch-clamp data is not widely accessible in large data sets because of its slow speed to acquire the data.

In order to further study the electrical properties, biophysical detailed models are created, thus allowing us to easily simulate many experiments in silico. These models try to recreate the behavior of real neurons by specifying the ion channels kinetics and their positions by constraining them with experimental morphological and electrical data (Druckmann et al., 2007; Gewaltig and Diesmann, 2007). This approach allows us to simulate experiments like injecting currents and recording them, even in dendrites and axon.

Once a biophysical model is created, we can create as many cell instances as we want, in order to study individual neurons or their connections in networks. However, if we want to create a realistic biophysical detailed model of a specific neuron, we need its morphological and electrical experimental data. The bottleneck therefore is the difficulty to acquire electrical data for many cells.

## 2 Objectives

To bypass this problem, here we propose a method to predict the electrophysiological data of single neurons based only on their morphology. A data set with both morphology and electrophysiology data for a set of neurons is provided to a supervised machine learning model. The model uses fully connected artificial neural networks to learn a multi-output regression method to predict multiple electrophysiological features taking multiple morphology features as the inputs.

As the data set contains only a small set of instances, the natural point estimates prediction of the neural network model could be imprecise. To further improve the model, uncertainty has been included so each output of the model represents a Gaussian distribution instead of just a point estimate.

The predicted outputs could be used for different research studies but the use case we present here is intended to be an aid for creating better biophysical detailed models of specific single neurons when we only have their morphology but not their electrical recordings, which is a very common scenario. The NeuroMorpho.org database stores more than 100.000 morphologies from different labs, which is a clear indicator of the massive morphology data available to create biophysical models.

To test the predictions of the model, a new morphology is given and the model outputs the predicted electrophysiological features. The morphology and the predicted electrical features are then provided to the BluePyOpt software (Van Geit et al., 2016) to create a biophysical detailed model to run in silico simulations.

Finally, we also provide an integration to run the obtained biophysical model within the Geppetto UI software (Cantarelli et al., 2018), which has the NEURON simulator (Hines and Carnevale, 1997) as its backend. All the software is written in Python and it is designed in a modular way to be able to include it in the NeuroSuites platform in the future.

## 3 Data set preprocessing and features

### 3.1 Data set

To learn this supervised model we need a data set with both morphology and electrophysiology data for each neuron in the data set. While large morphology databases are freely available, it is not the same with electrophysiology data. There exist platforms like NeuroElectro (Tripathy et al., 2014) that provide access to electrical data but the majority of the linked data lack the morphology so they are not valid for this study.

As far as we know, the only open data set with both morphology and electrophysiology data for the same neurons, is the Allen Cell Types database (Gouwens et al., 2018). However, in this data set we still have the problem of the low spatial precision for the electrophysiology. The patch-clamp is only done in the soma, therefore we do not have information about how the neuron would react to stimuli in the dendrites or axon. In other words, it would actually be impossible to create a model that can associate multiple parts of the morphology with the electrophysiology since we only have stimuli and recordings for the soma.

The native solution to this problem would be a data set with morphology and electrophysiology data from the soma, dendrites and axon. This means injecting currents in many different parts of the cell and recording the activity in many parts as well. This way we would have a rich set of features associating the morphology with the electrical behavior.

Since there are no data sets of this kind, another alternative is to use data from literature and expert knowledge to build biophysical detailed models with the morphology data and patch-clamp data in the soma. Once the biophysical models are built, we can run any kind of experiment in silico and simulate patch-clamp recordings in as many parts as we want, so we can obtain their electrical features. This procedure requires advanced fitting methods to be able to replicate the observed data for every specific neuron to be modeled. While the native solution is more precise, this last approach can also solve the problem in a efficient way because the study of biophysical models is well established since decades and simulations can be very precise.

The Allen Cell Types data set contains 637 neurons with both morphology and electrophysiology data (as of February 2020) recorded in the same experimental conditions (see (Gouwens et al., 2018) for detailed information). However, only 104 have their corresponding all-active biophysical detailed models. Perisomatic models were not included as they do not include active conductances in the dendrites. These include pyramidal and aspiny neurons (e.g. Pvalb positive) neurons from the visual cortex of the mouse. The fact of including refined biophysical models already created, makes this data set the perfect candidate for this supervised machine learning task. The data set used in this paper can be obtained in https://celltypes.brain-map.org/data, selecting the checkbox for “All-active biophysical models” and in the reconstruction type section, selecting “full” and “dendrite only”.

This can seem like a contradiction to our original problem, since the approach of creating a biophysical model with previous knowledge can also obtain the electrical features without having the experimental recordings but using fitting procedures to check the validity of resulting models (Markram et al., 2015).

So why would someone prefer to use the predictive approach of this paper in opposite to building a biophysical model with data from literature and expert knowledge, which can then also be used to simulate experiments and obtain the electrical features? Creating a biophysical model requires previous knowledge (from experts and literature) about the specific kind of neurons that we are studying. Our approach in the other hand is a supervised machine learning method, which by definition learn from this previous knowledge to make predictions in new instances for which we do not have this previous knowledge.

Therefore, our method is extensible to all kind of neurons as it works in a more individualized way, every predicted feature for a specific neuron depends exactly on the morphology of that neuron, and not in a general property of a group of neurons that have been analyzed in the literature.

It is important to note that these two approaches are not mutually exclusive. In fact, the ideal use for our method is to have some experts in the research team to check and refine the predicted electrical features obtained with the model.

### 3.2 Feature extraction and selection

The morphological features were provided in the Allen Cell Types data set, which were extracted using the L-Measure tool (Scorcioni et al., 2008) (Table 2).

**Table 1.**
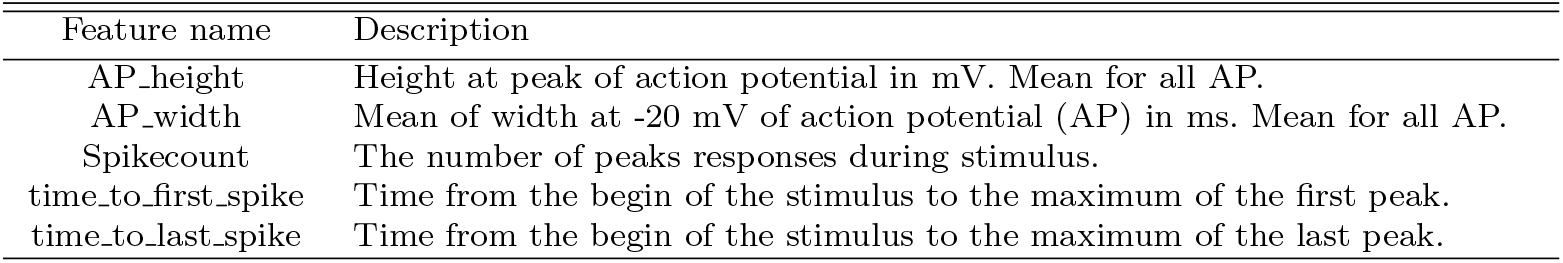
Electrophysiological features (information extracted from the Allen Cell Types database, biophysical models all active white paper (Allen Institute for Brain Science, 2016a))

**Table 2.** Morphological features (information extracted from the Allen Cell Types database, morphology white paper (Allen Institute for Brain Science, 2016b))

| Feature name | Description |
| --- | --- |
| average_diameter | The average diameter of all compartments of the neuron. |
| max_branch_order | The maximum order of the branch. A branch's order is defined with respect to the soma where the soma has a branch order = 0. The first bifurcation has a branch order = 1, the second bifurcation has a branch order = 2, and so on. |
| max_euclidean_distance | The maximum Euclidean distance of all nodes. Euclidean distance is the straight line distance from the soma (root) to the node. |
| max_path_distance | The maximum path distance of all nodes. The path distance is the sum of lengths of all connected nodes from the soma, ending with that node. |
| number_bifurcations | The number of bifurcations for the given input neuron. A bifurcation point has two daughters. |
| number_branches | The number of branches in the given input neuron. A branch is one or more compartments that lie between two branching points or between one branching point and a termination point. |
| number_nodes | The total number of nodes in the given input neuron (one SWC file). A node in a SWC file represents a single sample point of the neuron defined by its X, Y and Z coordinates, a radius, and its connectivity to other nodes in the neuron. |
| number_stems | The number of stems attached to the soma. When the type of the node is not soma, it is labeled as a stem. |
| number_tips | The number of terminal tips for the given input neuron. This function counts the number of nodes that are terminal endpoints. |
| overall_depth | Depth is measured on the y-coordinates and it is the difference of minimum and maximum y-values after eliminating the outer points on either ends by using the 95% approximation of the y-values of the given input neuron. |
| overall_height | Height is computed on the y-coordinates and it is the difference of minimum and maximum y-values after eliminating the outer points on either end by using the 95% approximation of the y-values of the given input neuron. |
| total_length | The total length of the neuron is computed as the sum of distances between two connected nodes for all branches of the input neuron. |
| total_surface | The surface of the soma is computed by one of two methods. If the soma is composed of just one node then the sphere assumption is used; otherwise the sum of the external cylindrical surfaces of nodes forming the soma is calculated. |
| total_volume | The total volume of the entire neuron. |

The Allen Cell Types data set provides the biophysical model files to run the simulations. We ran the biophysical models in the Python interface for the NEURON software (Hines et al., 2009) to extract the electrophysiological features. Our goal here is to run multiple simulations in order to inject and record the electrophysiology currents from different parts of the morphology.

The electrical features were therefore extracted from the simulations sweeps obtained by running the biophysical models in the NEURON software. The stimuli were long square current injections, i.e. a square pulse of 200 ms to allow the neuron to come to a steady state. The features of multiple sweeps corresponding to the responses of the cell to the currents stimuli were extracted using the eFEL software (Blue Brain Project, 2015) (Table 1, Fig. 2). This software outputs the necessary features to then feed to the BluePyOpt software to construct biophysical models. Additionally, some metadata features available in the dataset were also included in the model (Table 3). All this feature extraction process is depicted in Fig. 1.

**Table 3.**
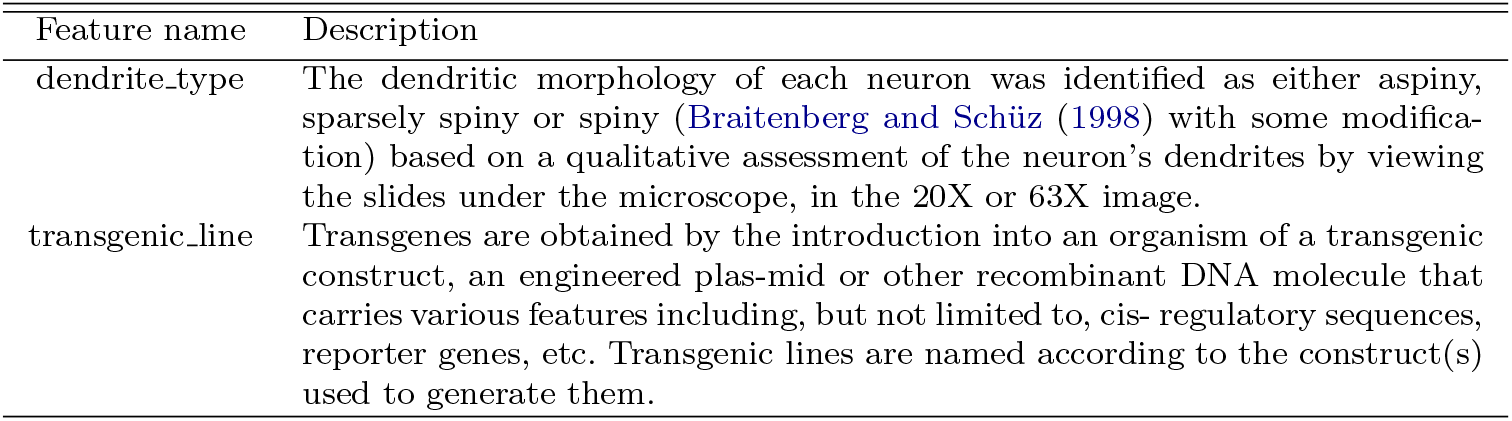
Metadata features (information extracted from the Allen Cell Types database, morphology white paper (Allen Institute for Brain Science, 2016b))

| Feature name | Description |
| --- | --- |
| dendrite_type | The dendritic morphology of each neuron was identified as either aspiny, sparsely spiny or spiny (Braitenberg and Schüz (1998) with some modification) based on a qualitative assessment of the neuron's dendrites by viewing the slides under the microscope, in the 20X or 63X image. |
| transgenic_line | Transgenes are obtained by the introduction into an organism of a transgenic construct, an engineered plas-mid or other recombinant DNA molecule that carries various features including, but not limited to, cis- regulatory sequences, reporter genes, etc. Transgenic lines are named according to the construct(s) used to generate them. |

**Fig. 1.**
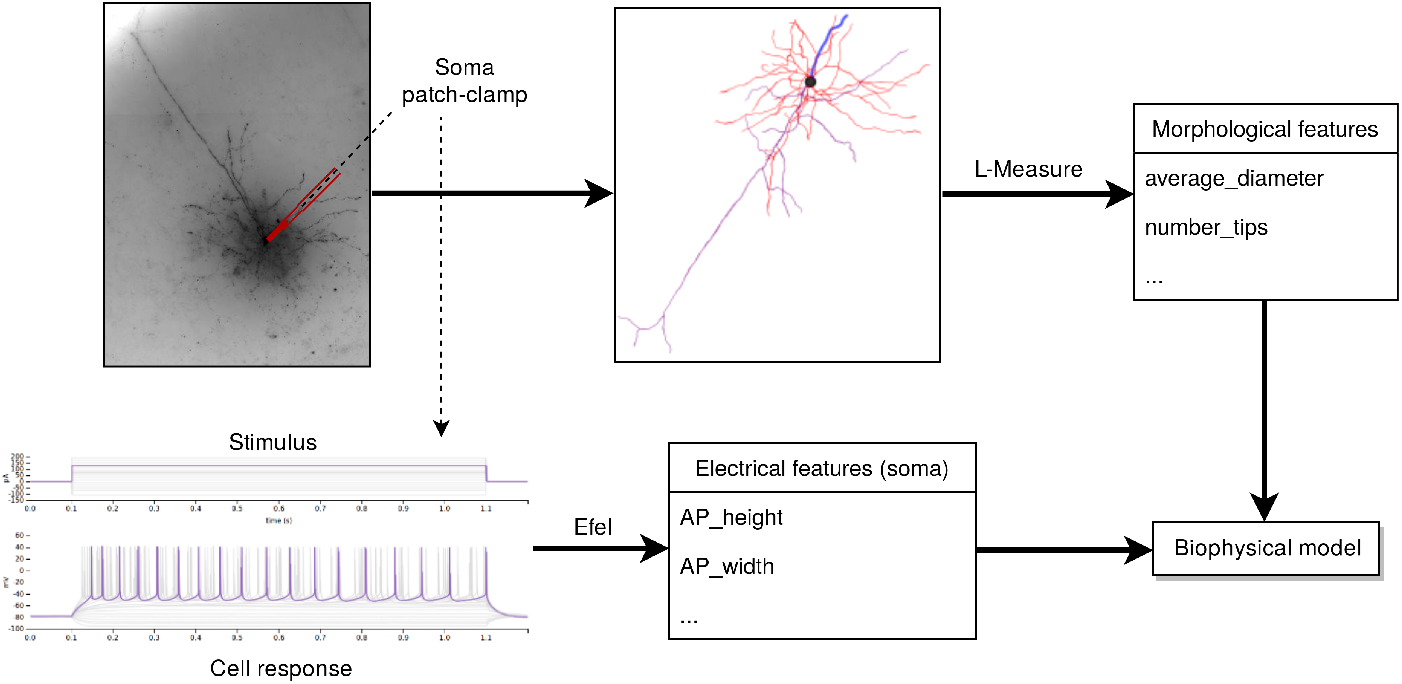
Data set features extraction. The morphology reconstruction is obtained from the microscopy image and The morphological features are extracted with the L-Measure software. Note that the morphology plot shows a 2D image for visualization purposes but 3D morphology features were also extracted as can be seen in Table 2. Electrophysiology sweeps data is obtained from the soma patch-clamp and the features are extracted with the eFEL software.

**Fig. 2.**
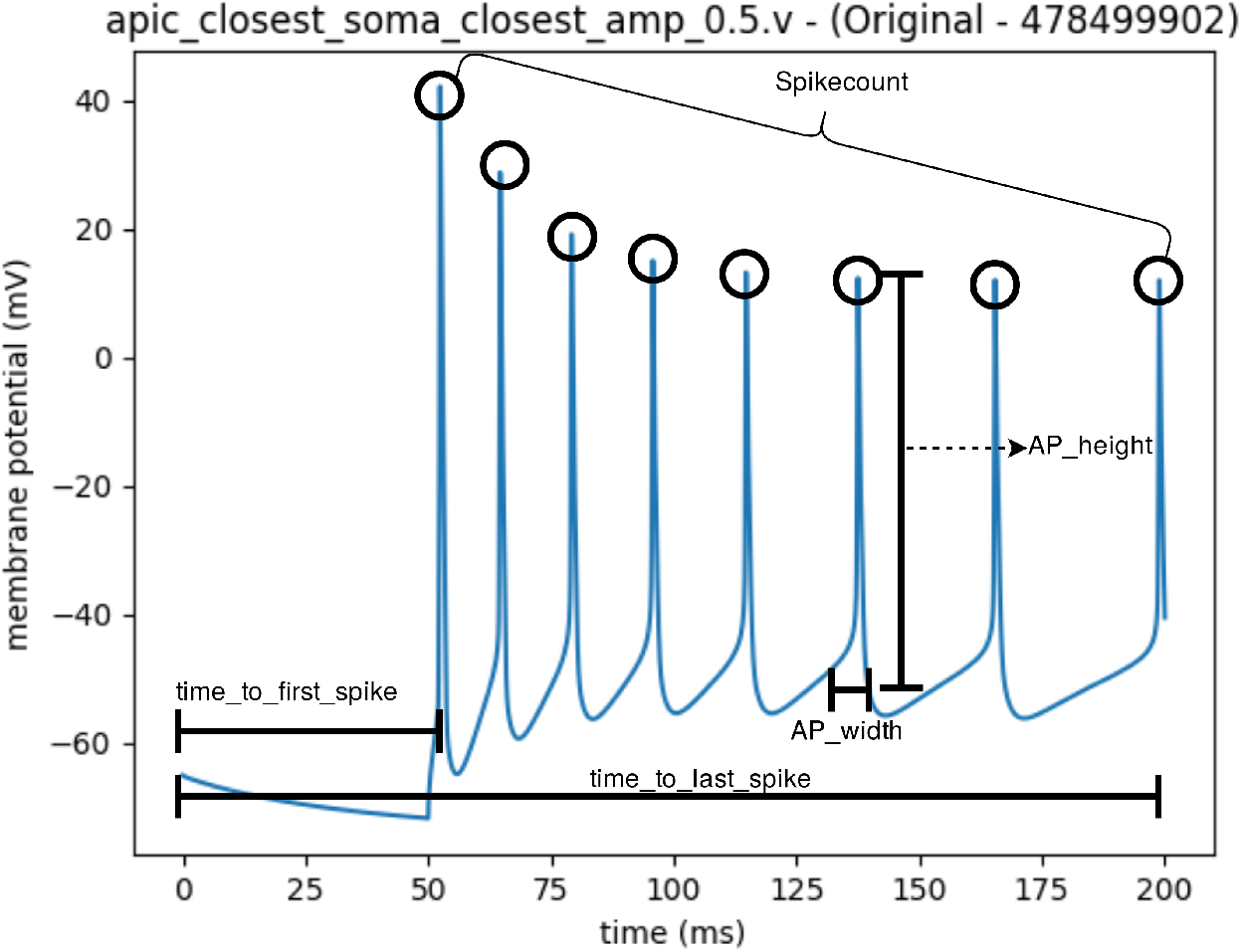
Electrophysiological features. Detailed description of the features is shown in table 1. Note that in this example the time_to_last_spike is almost at the end of the plot but this is just a coincidence for this precise example. Another example of an experimental trace can be seen in Figure 6.

To find the best morphological and electrical features, we selected those with highest Pearson correlation coefficient and mutual information between each pair of morphological-electrical feature. Mutual information is given more weight as it is capable of finding nonlinear relationships between the features.

We tested multiple electrophysiological sweeps configurations in a greedy search with the following variables: *iclamp section - record section - stimulus amplitude*, where *iclamp section* refers to the morphology part where the current was injected (soma, dendrite or axon), *record section* refers to the morphology part where the neuron response was recorded (soma, dendrite, axon), and *stimulus amplitude* refers to the input current amplitude (0.4 pA, 0.5 pA or 0.9 pA). Then, in order to find the best subset of configurations, we selected those with highest Pearson correlation coefficient mean and mutual information mean between all the features in the morphological-electrical features of each configuration (see Table 5 and 6 to check the best configurations obtained from this process).

**Table 4.** Best neural network hyperparameters

| Data set | Number of hidden layer | Neurons per hidden layer | Dropout rate | Learning rate | Number of epochs |
| --- | --- | --- | --- | --- | --- |
| Group A | 3 | 616 | 0.6 | 0.001 | 289 |
| Group B | 3 | 616 | 0.6 | 0.001 | 1128 |

**Table 5.**
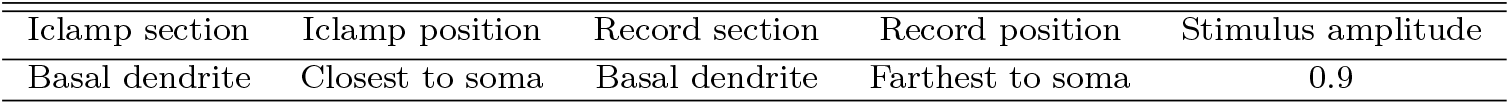
Group A electrical configurations

**Table 6.**
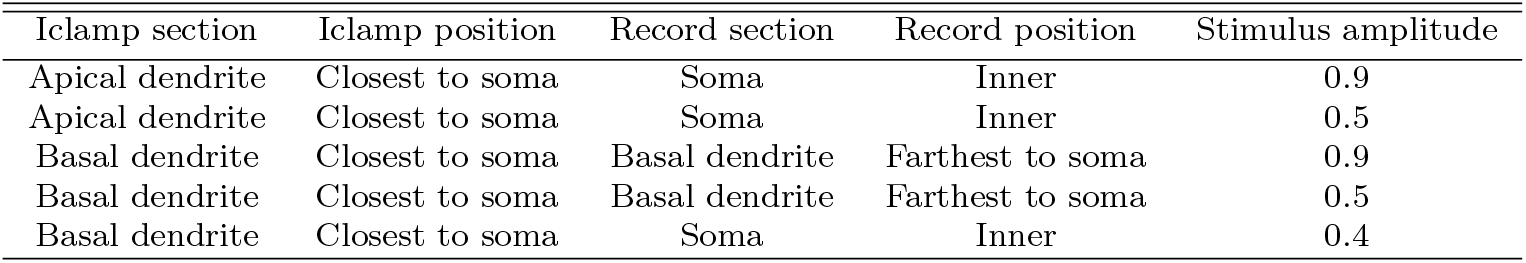
Group B electrical configurations

## 4 Machine learning

### 4.1 Multi-output regression with neural networks

We opted to learn a multi-output regression model in order to be able to predict specific electrophysiology outputs to specific morphology inputs, rather than grouping the neurons in classes with general properties to train a classifier.

The selected model is a fully connected neural network, as it allows to represent nonlinear relationships between the features and also take into account the relationships between the output features thanks to the back propagation algorithm. The morphological features correspond to the inputs and the electrical features obtained with the simulations of the biophysical models are used to compute the cost function (Fig. 3).

**Fig. 3.**
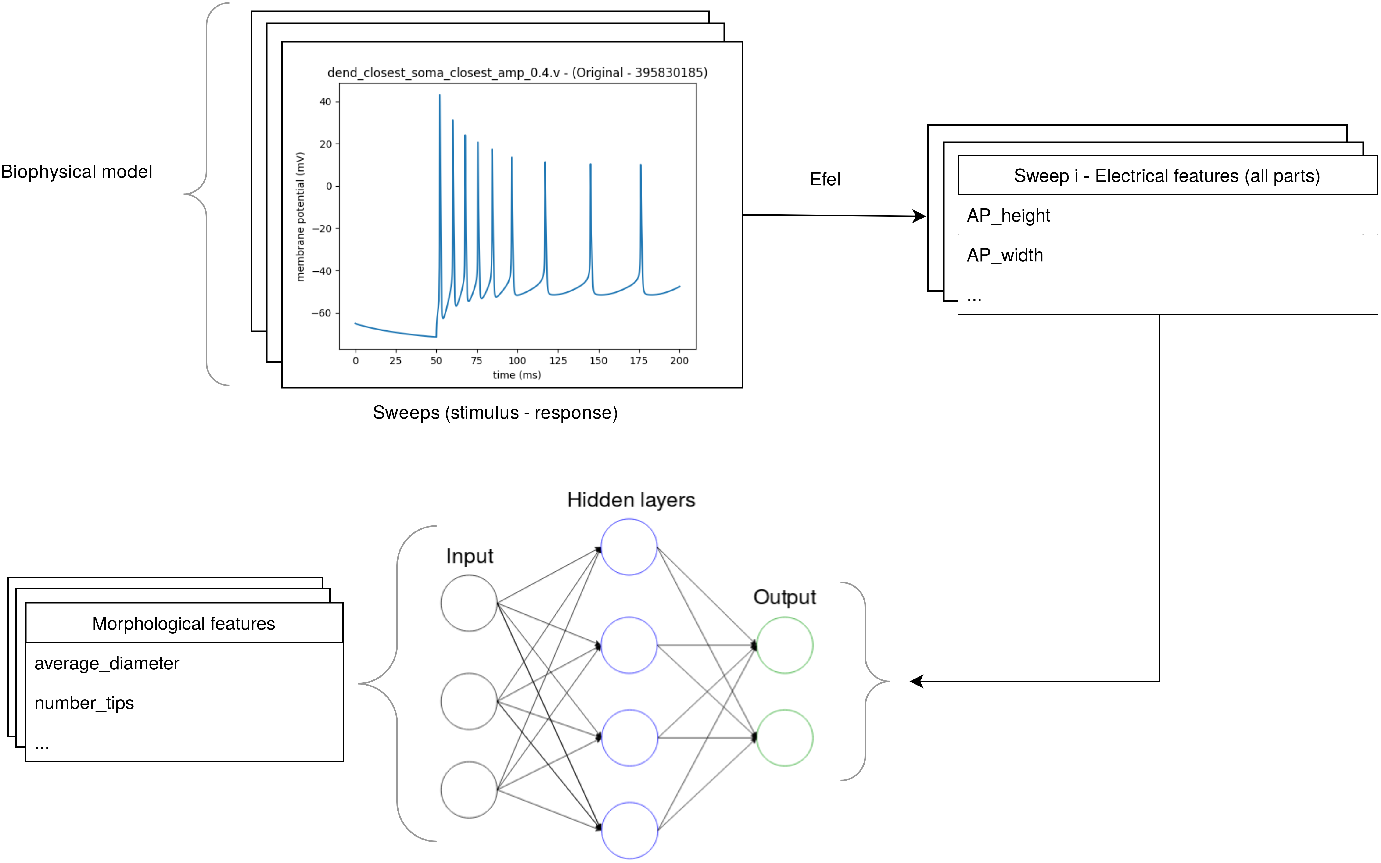
Machine learning supervised training. The neural network model is trained taking The morphological features as the inputs. To compute the cost function we use the electrophysiological features from the biophysical model sweeps of the same neurons.

### 4.2 Including uncertainty in neural networks

The data set contains very few instances for a model to predict precise outputs in unseen data. Furthermore, one drawback of neural networks is the lack of expressivity to account for uncertainty, as the results obtained in a regression problem are only point estimates.

When dealing with uncertainty we must differentiate two kinds of it. Aleatoric uncertainty happens when the data have some kind of noise in the input measurements that produce uncertain outputs. Aleatoric uncertainty can either be homoscedastic if the noise is constant for all the inputs or heteroscedastic if the noise varies for every input (Brando et al., 2019). This kind of uncertainty cannot be reduced even if the data set had more instances.

Epistemic uncertainty in the other hand refers to the ability of the model parameters to take into account the uncertainty in the model itself. Therefore one native way to reduce this uncertainty would be to have more instances.

In our data set both the inputs and the outputs are likely to have aleatoric heteroscedastic uncertainty since the measurementes are very dependent of the experimental conditions and some techniques like patch-clamp are so complex that are prone to small measurement errors. However, the most common design of neural networks is not able to produce uncertainty in the outputs prediction, as they only output point estimates as the regression result.

To take into account this aleatoric heteroscedastic uncertainty, we replaced the *n* output features of the final layer with 2*n* outputs, so for each output feature we now have its mean and standard deviation of a normal distribution (Fig. 4):

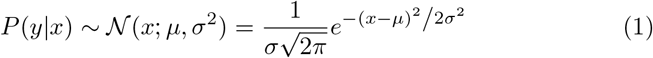

**Fig. 4.**
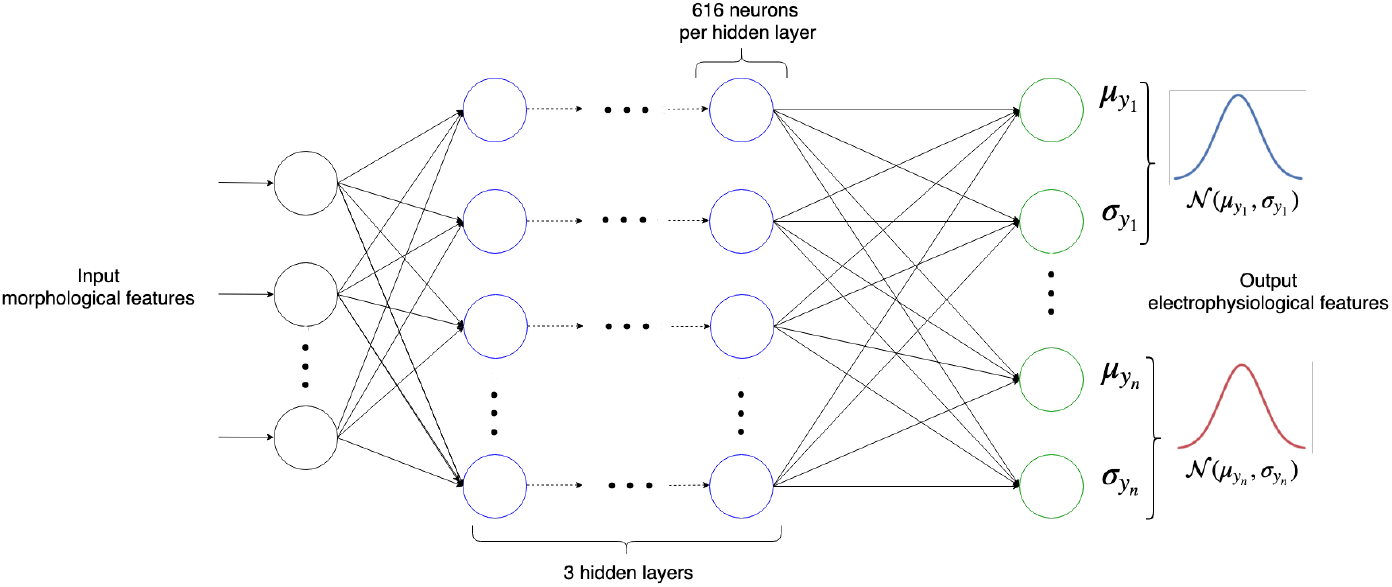
Neural network design. Hyperparameters are obtained via grid-search optimization (table 4). Output features consist in pairs of *μ*_*i*_, *σ*_*i*_ to create a normal distribution for each of the electrophysiological features to be predicted. Negative log-likelihood (equation 4) is used as the cost function.

This model can be seen as a simple version of the Mixture Density Model (MDN) (Bishop, 1994), where a mixture of Gaussians are placed for every output feature instead of a single Gaussian

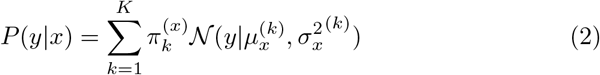

For completeness we also tested a MDN model but it did not improve the performance over the single Gaussian model so we stuck with the simplest model for computational efficiency.

To train the model, the mean squared error (MSE) (equation 3) loss (i.e. cost function) is not valid anymore since now we must calculate the maximum likelihood estimation (MLE) of the Gaussian distribution for each output feature given the inputs.

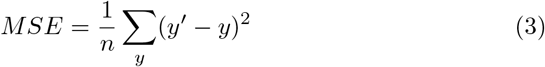

where *y* are the ground-truth values and *y′* the predicted values for the instances.

Therefore the negative log-likelihood loss (NLL) (equation 4) is used as a cost function. The negative sign is just a computational requirement because the gradient descent optimization algorithm for neural networks minimizes the objective error.

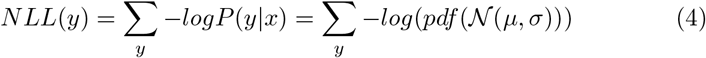

To capture the epistemic uncertainty, we included a dropout factor after each activation function in every layer, as dropout can be interpreted as a variational approximation (Gal and Ghahramani, 2016). This also allows us to better generalize the model as it works as a way of regularization.

### 4.3 Hyperparameters selection

The model hyperparameters are specially important in this case as we have very few instances and multiple features as outputs. A grid search was executed, and for every configuration of hyperparameters, the best values obtained in a 4 k-fold cross validation were selected (Table 4). The model was implemented in Python using the Pytorch library (Paszke et al., 2017).

## 5 Results

Neuronal morphologies were already processed by the Allen Institute by applying a reconstruction repair algorithm for the cases when some dendrites or axon were cut in the plane when doing the reconstruction.

However, some neurons in the data set do not have a complete morphology reconstruction, i.e. the apical dendrites are missing. This presents a problem when extracting the new electrical features with the biophysical model simulation, since we can not inject or record currents in the apical dendrites.

To take this into consideration, we made two copies of the data set: one containing only the electrical features corresponding to the injections/recordings where the apical dendrites were not involved (group A: Table 5), and another one containing all kind of electrical features (group B: Table 6). Note that both groups contain all the instances, the differences are only the electrical features. Therefore, in the group B, the neurons without apical dendrites in the reconstruction will have zeros in those predictions involving currents in the apical dendrites, since the simulation would be impossible.

Lastly, we split the data set in training (80% of the total number of instances) and test (20% of the total number of instances) sets. In the training set we selected the best hyperparameters for the model using the k-fold cross validation and once we had selected the best hyperparameters we ran a last training over all the training set. The test set was never used to train the model as it is used to check how the model performs in unseen data.

### 5.1 Metrics (MSE, NLL)

To provide a benchmark, we compared the results with other known regression methods like regularized linear regression and random forest regressor, already implemented in Scikit-Learn (Pedregosa et al., 2011). Furthermore, we included the untrained neural network model in the comparison to understand how is performing the trained model compared to an untrained one. Note however than the untrained neural network will not have such bad performance as a random model because the untrained model makes use of advanced parameters initialization methods like the common known as Xavier initialization method (Glorot and Bengio, 2010).

To have a fair comparison, we also ran a cross validation grid search over the linear regression method and the random forest regressor to find their best hyperparameters. However, all these model give us point estimates instead of the probability distribution output that our model predicts. Hence to make comparisons between these models we had to calculate the MSE. The exception here is the untrained neural network model, where we do have a probability distribution output so we can compare it to the NLL of our model.

To calculate the MSE with our prediction, we took the mean of our Gaussian probability distributions, while the uncertainty (*σ* in our case) is included in the NLL formula.

Group A results are in Table 7 and Fig. 5. Group B results are in Table 8.

**Table 7.** Group A results

| Method | MSE train set | MSE test set | NLL train set | NLL test set |
| --- | --- | --- | --- | --- |
| Linear regression | 1744.88 | 4070.00 | - | - |
| Ridge linear regression | 1918.41 | 2789.74 | - | - |
| Lasso linear regression | 1993.36 | 2687.1 | - | - |
| Random forest | 1611.89 | 2495.25 | - | - |
| Untrained neural network | 2138.69 | 2627.27 | 1.96 | 1.96 |
| Neural network | **1049.96** | **1751.05** | **1.48** | **1.83** |

**Table 8.** Group B results

| Method | MSE train set | MSE test set | NLL train set | NLL test set |
| --- | --- | --- | --- | --- |
| Linear regression | 1076.39 | 2165.50 | - | - |
| Ridge linear regression | 1241.86 | 1706.44 | - | - |
| Lasso linear regression | 1269.46 | 1547.50 | - | - |
| Random forest | 1078.09 | 1371.56 | - | - |
| Untrained neural network | 2282.45 | 2509.277 | 3.57 | 3.57 |
| Neural network | **577.89** | **1091.29** | **2.16** | **3.31** |

**Fig. 5.**
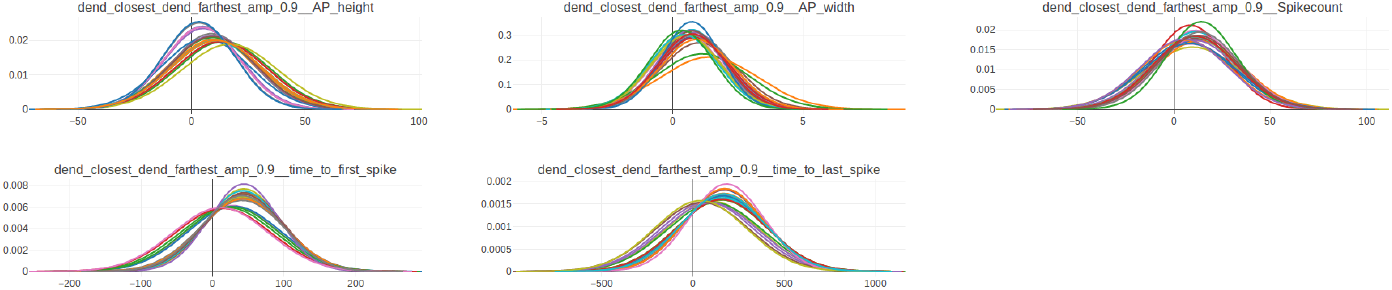
Group A results in the test set for dend_closest_dend_farthest_amp_0.9 sweep. Each chart represents the predictions of an electrical feature. Each normal distribution in every chart corresponds to the predicted feature of a neuron. For instance, the first chart represents the AP_height predicted feature for every neuron in the test set (21 neurons, i.e. 20% of total instances in the data set), so each normal distribution (each one in a different color) corresponds to the predicted AP_height for that neuron in this specific sweep.

To showcase a specific example, we selected a random neuron in the test set to run the model inference. Then we compared the predicted results to the ground truth data (Fig. 6).

**Fig. 6.**
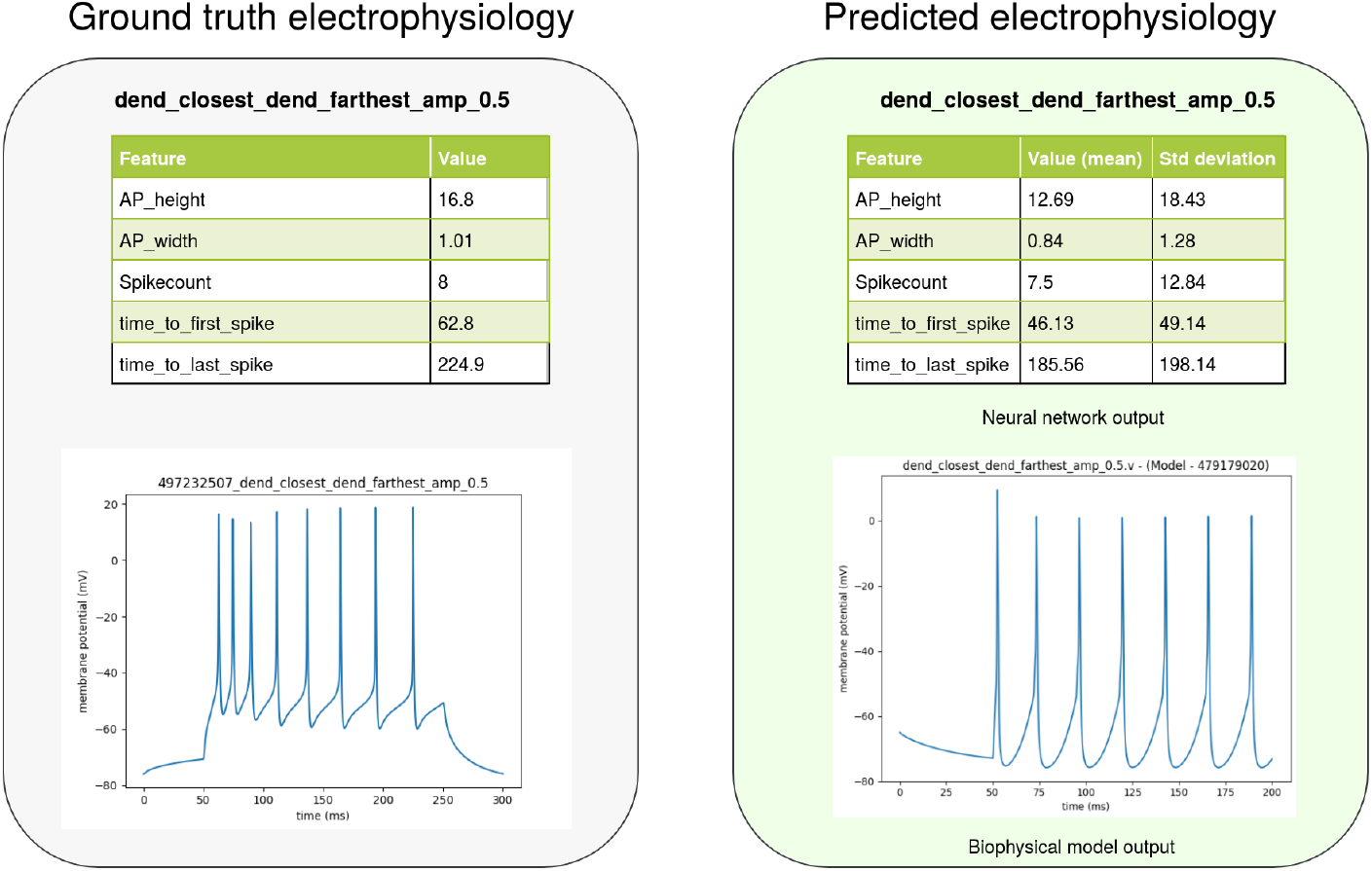
Example of two electrophysiology predicted sweeps. Use case example for an out-of-sample neuron in the test set in the Allen Cell Types data set. Left: ground truth electrophysiological features obtained as shown in Fig. 1. Right: predicted electrophysiological features for the two sweeps and the corresponding biophysical model optimized by BluePyOpt. Note that the biophysical model building does not match exactly with the neural network output since biophysical models try to recreate experimental conditions but they are not perfect. Even if the electrical features were identically the same as the ground truth, the biophysical model trace would differ a little since they are models after all. However, the focus of this paper is the predictions of the electrophysiology and therefore the neural network output are used to generate the results. The biophysical model building is just an use case of the method to illustrate the results in a visual way. Ids associated in the Allen Cell Types data set: Specimen id: 479179020 Biophysical model id: 497232507

### 5.2 Biophysical models simulation and use case

With the morphology reconstruction and the eletrophysiology features we can run optimization algorithms to set the most precise configuration of ion channels to build a biophysical model. We leveraged this task to the BluePyOpt tool, which also provides functionality to set a standard deviation for the optimization of each feature. Finally, we developed the necessary transformations for the obtained biophysical model to make it compatible to the Geppetto UI platform, which uses the NEURON software as the backend to run the biophysical model. Using the Geppetto UI software allows us to run simulations in a sophisticated user interface to be able to get great insights. To provide a production ready software, we created a software pipeline to connect all these processes (Fig. 7).

**Fig. 7.**
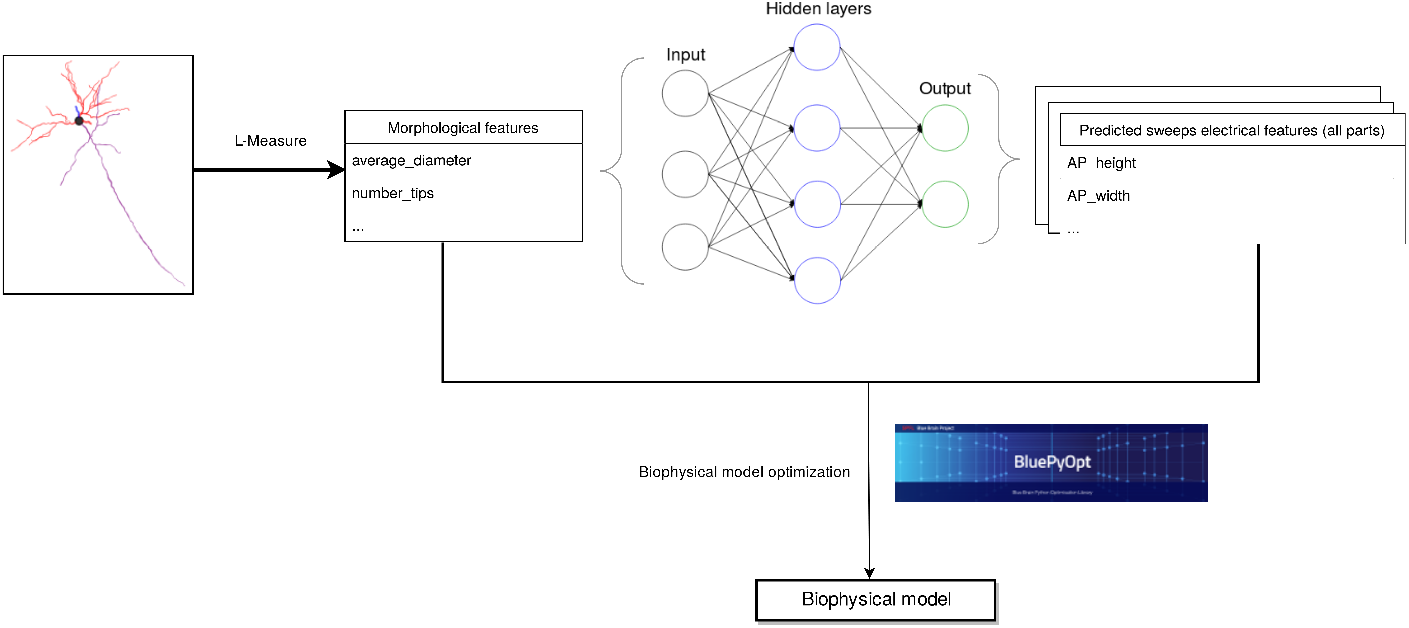
Machine learning inference in new data. Once the model has been learned, we use it to predict the electrophysiological features of neurons where we only have their morphology data.

As we said before, the data set has a very small number of instances, meaning that the prediction may not be completely accurate. In order to build accurate biophysical models, it is desirable to have background experience in the field to be able to verify the predictions and adjust them, so the predicted features would serve as support to build better models.

To present a use case, we downloaded the morphology of a set of pyramidal neurons with complete morphology reconstructions from NeuroMorpho.org. The desired neuron reconstructions were downloaded using our user interface in NeuroSuites. This way we can also run the L-Measure tool online in these neurons and finally download the morphology reconstructions in swc format (Cannon et al., 1998). One neuron was randomly selected by searching pyramidal neurons with complete morphology.

Once the model outputs the predicted electrical features, all we have to do is build the biophysical model with BluePyOpt (the final result can be seen in Fig. 8) and then exporting the model to Geppetto (Fig. 9) where we can run all kind of simulations.

**Fig. 8.**
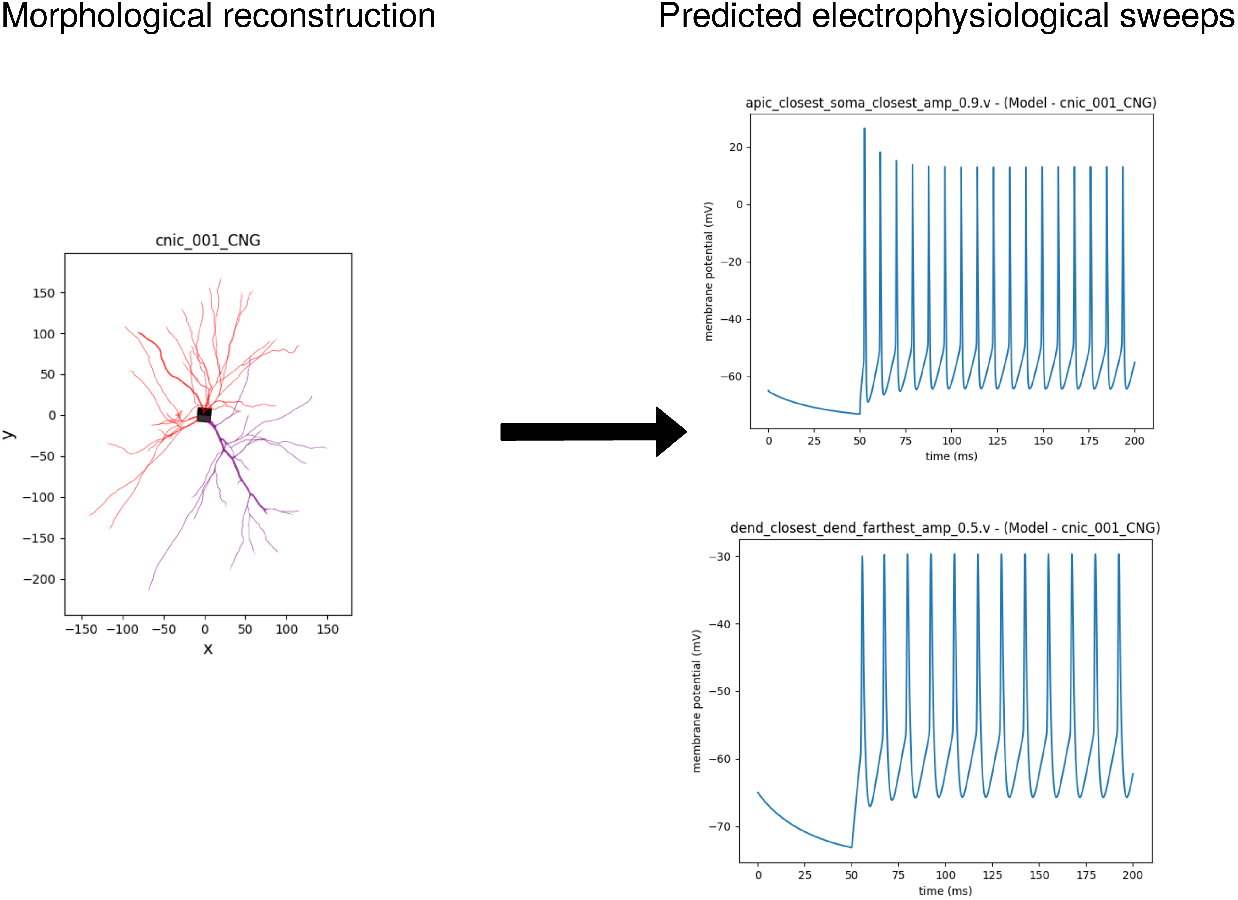
Electrophysiology sweeps predicted in an out-of sample neuron from NeuroMorpho.org. Sweep plots were created using the BluePyOpt biophysical model optimization generated from the predicted electrophysiology data. Specimen id associated in NeuroMorpho.org: CNIC_001

**Fig. 9.**
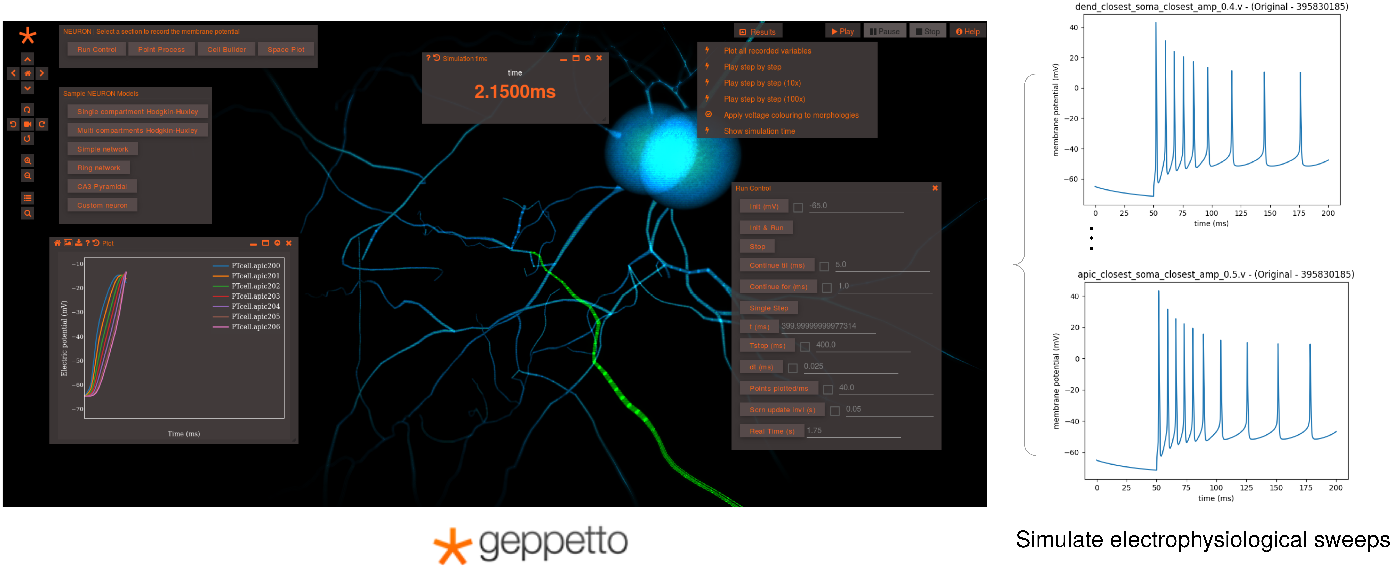
Geppetto UI. Biophysical model obtained by the BluePyOpt optimization from the predicted electrophysiology data, running in the Geppeto UI and generating new patch-clamp simulations in real time.

Depending on the data set, some morphology and metadata features from the original Allen Cell Types data set may not be available in different data sets from other sources. Morphology features can be extracted from the original data, however, the original data may not be available and only the features would be provided. Metadata features, however, must be provided in the data set. As NeuroMorpho is the largest morphology reconstruction database, we included a pretrained model that contains only the features available in its database. This pretrained model without specific metadata allows for a better translation to different datasets. The only difference in this one is removing the missing features in the NeuroMorpho features with respect to the Allen Cell Types data set features, i.e. *number nodes*, *dendrite type* and *transgenic line*.

## 6 Discussion

Biophysical models are a key element for simulating the electrical properties of the neurons but the lack of large databases of experimental data to create these realistic models has always been a problem for the ion channels configuration.

This method presents a starting point for these kind of predictions. The multiple drawbacks coming from the raw data (small number of instances, patch-clamp only in the soma, etc) makes this method not recommended to blindly trust in its predictions as the only tool for building the biophysical models. Then, the proper use is as an aid for the experts setting the ion channels configuration for building the biophysical models. The uncertainty in the output prediction allows the humans to have a clearer perspective of the range of most probable values for every electrical property. These insights plus the expert knowledge can improve the quality of the biophysical models.

A possible improvement to the model would be the availability of more heterogeneous data sets. The Allen Cell Types neurons containing electrophysiology, morphology and their corresponding all-active biophysical model are very homogeneous, meaning that we have data from only one species (mouse), brain region (visual cortex) and two morphology types (pyramidal and aspiny neurons). The only heterogeneous metadata in this data set are the dendrite type (spiny, aspiny, sparsely spiny) and transgenic line. Hence, if new data sets also include more heterogeneous instances, they would contribute to having a richer metadata features like different species, brain regions, morphology types, etc.

Research in electrophysiology acquisition methods is greatly advancing in automation tasks and now some methods are able to capture the electrical properties many neurons in faster ways (Holst et al., 2019; Jouhanneau and Poulet, 2019). In the future is expected that larger data sets will be published and the accuracy of this method could be greatly improved.

On the other hand, we modelled the epistemic uncertainty including a dropout factor but there are also alternative machine learning models exist to deal with it like Bayesian neural networks (Blundell et al., 2015), which assign a probability distribution for every weight in the neural network instead of just point estimates.

Base template models are used for building the final biophysical model with the BluePyOpt software. This allow us to include expert knowledge about specific neuron types that can greatly improve the quality of the model. The base model used in this paper is a layer 5 pyramidal neuron from the neocortex (Markram et al., 2015), which can be valid for the data set we used as the neurons are also pyramidal. The availability of different base models would allow us to better fit the parameters of neurons where we have useful expert knowledge about their type.

Finally, in the future we aim to include this software in our NeuroSuites platform. This will allow the users to take advantage of the many tools offered in NeuroSuites and the option to interconnect them with this software. The idea would be to create a workflow where we would be able to run everything in NeuroSuites. The first step would be selecting the neuron morphologies from the NeuroMorpho filtering page and then extracting their features with the L-Measure tool also available in our platform. The extension would be to integrate the rest of the workflow in NeuroSuites, thus applying our pretrained models in order to obtain their predicted electrophysiological features. Finally the biophysical model would be created by the BluePyOpt optimization and visualized with the Geppetto UI software.

## Information Sharing Statement

All the software source code, data sets and instructions to run the software are freely available in our Gitlab public repository: https://gitlab.com/mmichiels/multi_output_regression_tfm.

## Acknowledgments

The authors declare no acknowledgments to cite.

